# Functional and Highly Crosslinkable HIV-1 Envelope Glycoproteins Enriched in a Pretriggered Conformation

**DOI:** 10.1101/2021.09.27.462085

**Authors:** Hanh T. Nguyen, Alessandra Qualizza, Saumya Anang, Meiqing Zhao, Shitao Zou, Rong Zhou, Qian Wang, Shijian Zhang, Ashlesha Deshpande, Haitao Ding, Amos B. Smith, John C. Kappes, Joseph G. Sodroski

## Abstract

Binding to the receptor, CD4, drives the pretriggered, “closed” (State-1) conformation of the human immunodeficiency virus (HIV-1) envelope glycoprotein (Env) trimer into more “open” conformations (States 2 and 3). Broadly neutralizing antibodies, which are elicited inefficiently, mostly recognize the State-1 Env conformation, whereas the more commonly elicited poorly neutralizing antibodies recognize States 2/3. HIV-1 Env metastability has created challenges for defining the State-1 structure and developing immunogens mimicking this labile conformation. The availability of functional State-1 Envs that can be efficiently crosslinked at lysine and/or acidic amino acid residues might assist these endeavors. To that end, we modified HIV-1_AD8_ Env, which exhibits an intermediate level of triggerability by CD4. We introduced lysine/acidic residues at positions that exhibit such polymorphisms in natural HIV-1 strains. Env changes that were tolerated with respect to gp120-gp41 processing, subunit association and virus entry were further combined. Two common polymorphisms, Q114E and Q567K, as well as a known variant, A582T, additively rendered pseudoviruses resistant to cold, soluble CD4 and a CD4-mimetic compound, phenotypes indicative of stabilization of the pretriggered State-1 Env conformation. Combining these changes resulted in two lysine-rich HIV-1_AD8_ Env variants (E.2 and AE.2) with neutralization- and cold-resistant phenotypes comparable to those of natural, less triggerable Tier 2/3 HIV-1 isolates. Compared with these and the parental Envs, the E.2 and AE.2 Envs were cleaved more efficiently and exhibited stronger gp120-trimer association in detergent lysates. These highly crosslinkable Envs enriched in a pretriggered conformation should assist characterization of the structure and immunogenicity of this labile state.

**IMPORTANCE:** The development of an efficient vaccine is critical for combating HIV-1 infection worldwide. However, the instability of the pretriggered shape (State 1) of the viral envelope glycoprotein (Env) makes it difficult to raise neutralizing antibodies against HIV-1. Here, by introducing multiple changes in Env, we derived two HIV-1 Env variants that are enriched in State 1 and can be efficiently crosslinked to maintain this shape. These Env complexes are more stable in detergent, assisting their purification. Thus, our study provides a path to a better characterization of the native pretriggered Env, which should assist vaccine development.

## INTRODUCTION

Human immunodeficiency virus type 1 (HIV-1) entry into target cells is mediated by the viral envelope glycoprotein (Env) trimer (1, 2). The Env trimer is composed of three gp120 exterior subunits and three gp41 transmembrane subunits (2). In infected cells, Env is first synthesized as an uncleaved precursor in the rough endoplasmic reticulum (ER), where signal peptide cleavage, folding, trimerization, and the addition of high-mannose glycans take place (3–6). Exiting the ER, the trimeric gp160 Env precursor follows two pathways to the cell surface (7). In the conventional secretory pathway, the Env precursor transits through the Golgi compartment, where it is cleaved into gp120 and gp41 subunits and is further modified by the addition of complex sugars. These mature Envs are transported to the cell surface and are incorporated into virions (8–11). In the second pathway, the gp160 precursor bypasses processing in the Golgi and traffics directly to the cell surface; these Golgi-bypassed gp160 Envs are excluded from virions (7).

Single-molecule fluorescence resonance energy transfer (smFRET) experiments indicate that, on virus particles, the mature (cleaved) Env trimer exists in three conformational states (States 1-3) (12). From its pretriggered conformation (State 1), the metastable Env trimer interacts with the receptors, CD4 and CCR5 or CXCR4, and undergoes transitions to lower-energy states (2, 12–23). Initially, the engagement with CD4 induces an asymmetric intermediate Env conformation (State 2) (24, 25). Binding of additional CD4 molecules to the Env trimer then induces the full CD4-bound, prehairpin intermediate conformation (State 3) (24–31). An extended coiled coil consisting of the heptad repeat (HR1) region of gp41 is exposed in the prehairpin intermediate (23, 25, 29–31). State 3 Env subsequently interacts with the CCR5 or CXCR4 coreceptor to trigger the formation of a gp41 six-helix bundle, a process that results in fusion of the viral and target cell membranes (32–36).

Env strain variability, heavy glycosylation and conformational flexibility contribute to HIV-1 persistence by avoiding the binding of potentially neutralizing antibodies. Mature Envs from primary HIV-1 strains largely reside in a State-1 conformation, which resists the binding of most antibodies elicited during natural infection (12, 23, 37–39); these high-titer, poorly neutralizing antibodies often recognize State-2/3 Env conformations (40–44). After years of infection, a small percentage of HIV-1-infected individuals generate broadly neutralizing antibodies (bNAbs), most of which recognize the State-1 Env conformation (12, 37, 38, 45–54). Passively administered monoclonal bNAbs have been shown to be protective in animal models of HIV-1 infection, suggesting that the elicitation of bNAbs is an important goal for vaccines (55–60).

Unfortunately, bNAbs have not been efficiently and consistently elicited in animals immunized with current HIV-1 vaccine candidates, including stabilized soluble gp140 (sgp140) SOSIP.664 trimers (61–69). Compared with functional membrane Envs, differences in the antigenicity, glycosylation and conformation of sgp140 SOSIP.664 trimers have been observed (70–77), potentially contributing to the inefficiency of bNAb elicitation. Single-molecule FRET (smFRET) analysis indicates that the sgp140 SOSIP.664 trimers assume a State-2-like conformation (78). These studies imply that the available structures of sgp140 SOSIP.664 and other detergent-solubilized Env trimer preparations (27, 28, 77, 79–91) differ from that of State-1 Env. The extent of the structural differences between the State-1 and State-2 Env conformations and their potential impact on Env immunogenicity are unknown. However, the importance of the State-1 Env as a likely target of vaccine-induced bNAbs provides a rationale for better characterization of this conformation.

HIV-1 is a polymorphic virus with a high mutation rate, allowing escape from host immune responses and antiretroviral drugs (92–99). Env polymorphisms that arise naturally or as a result of tissue-culture adaptation can result in altered virus infectivity, receptor binding or neutralization sensitivity (23, 40, 44–46, 100–119). Specifically, changes in “restraining residues” in gp120 have been shown to destabilize State 1, disrupt the closed pretriggered Env conformation, and lead to increased sampling of downstream conformations (45, 118, 119). These more “triggerable” Env mutants exhibit increased sensitivity to cold, soluble CD4 (sCD4), CD4-mimetic compounds and poorly neutralizing antibodies (23, 37, 45, 118, 119). Less common Env alterations apparently decrease Env triggerability and stabilize a State-1 conformation (120–125).

Crosslinking of HIV-1 Env amino acid residues, in some cases combined with mass spectrometry, has been used to study Env conformations (37, 73, 126–131). Crosslinking protocols that target lysine or acidic amino acid residues on native proteins have been integrated with mass spectrometry to provide low-resolution structural information (132–135). Here, we introduced lysine and acidic amino acid residues into a primary HIV-1 Env, using natural polymorphisms as a guide. Env changes that were functionally tolerated were combined to create Envs that are potentially able to be conformationally fixed by treatment with specific crosslinking agents. In the process of generating these Env variants, we identified two common polymorphisms that increased virus resistance to cold, sCD4 and a CD4-mimetic compound, phenotypes associated with stabilization of a pretriggered (State-1) Env conformation (120–125). Two lysine-rich variants with cold- and sCD4-resistant phenotypes were cleaved more efficiently and exhibited stronger gp120-trimer association in detergent lysates compared with the parental HIV-1 Env. Such highly crosslinkable Envs enriched in a pretriggered conformation should assist characterization of State 1.

## RESULTS

### Env variants with common lysine and acidic residue polymorphisms

We sought to create functional primary HIV-1 Env variants with an increased number of lysine/acidic residues that could be used to introduce stabilizing crosslinks. To identify Env residues that might potentially tolerate such substitutions, we compared Env sequences from 193 Group M, N, O and P HIV-1 and SIV_cpz_ strains (136). We identified Env residues where lysine or acidic substitutions occurred in at least 5% of these natural virus strains from more than one phylogenetic clade. The lysine polymorphisms were grouped by location in Env regions (gp41 and gp120 C-terminus, gp120 trimer association domain and gp120 inner domain) and by the number of substitutions in a set (Sets 4-7 contain additional lysine substitutions compared with those in Sets 1-3) (Fig. 1A). The ED2 set contains seven of the most common aspartic acid and glutamic acid polymorphisms in natural HIV-1/SIV_cpz_ variants (Fig. 1A).

**Figure 1.**
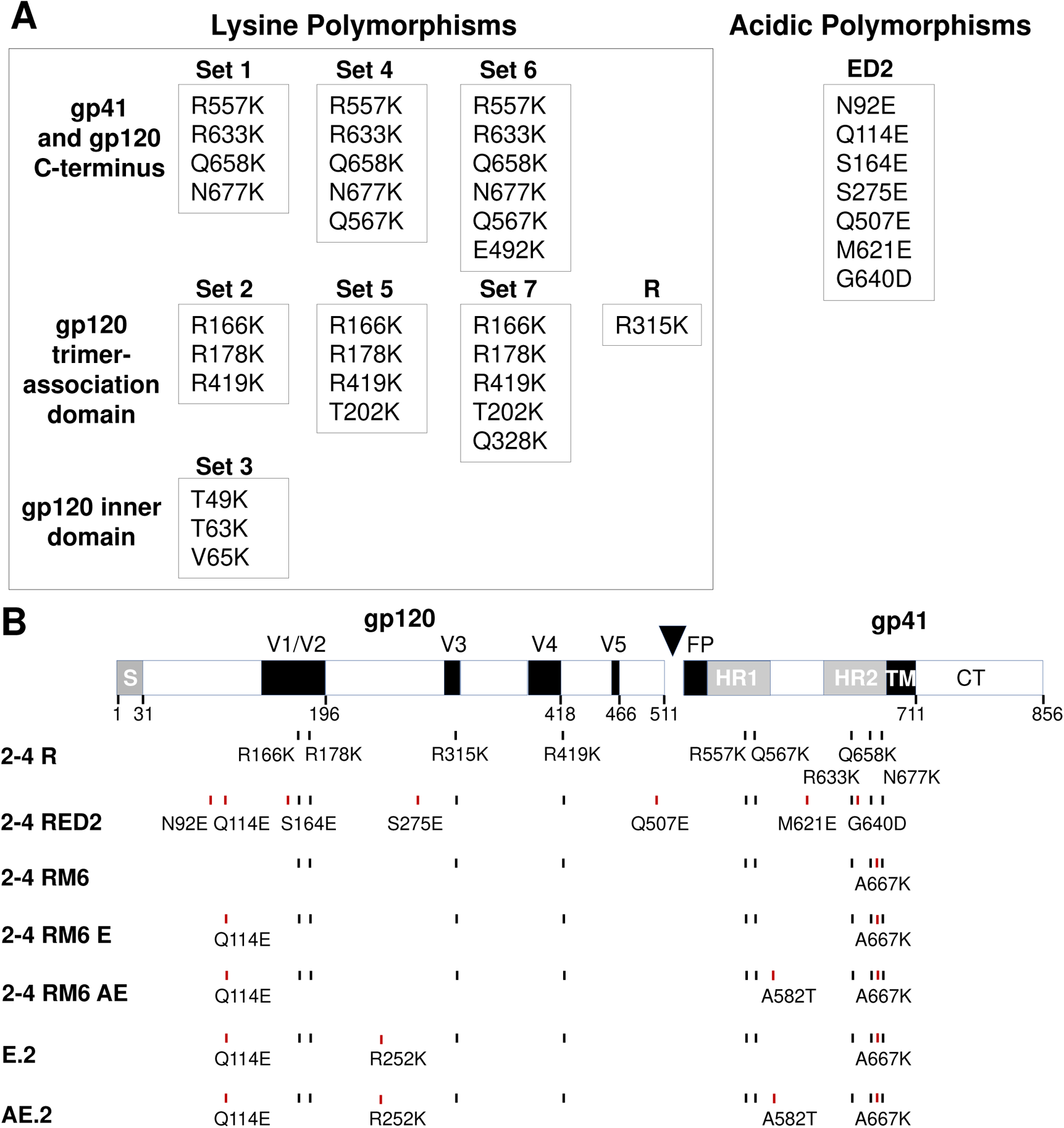
HIV-1_AD8_ Env modification guided by natural polymorphisms. (A) Natural polymorphisms in HIV-1 Env were used to suggest amino acid residues that might tolerate replacement with a lysine residue or with acidic amino acid residues. The lysine substitutions are grouped according to the Env region in which the residues are located. Compared with Sets 1-3, Sets 4-7 contain an increased number of substitutions. (B) A schematic representation of the HIV-1_AD8_ Env glycoprotein is shown, with the gp120-gp41 cleavage site depicted as a black triangle. S, signal peptide; V1-V5, gp120 major hypervariable regions; FP, fusion peptide; HR, heptad repeat region; TM, transmembrane region; CT, cytoplasmic tail. The amino acid changes associated with some of the key Env variants studied here are shown. Red vertical tick marks indicate changes in addition to those found in the 2-4 R Env.

We selected the primary Clade B HIV-1_AD8_ as the source of the parental “wild-type” Env in this study. Primary HIV-1 Envs differ in triggerability by CD4, a property that influences virus resistance to sCD4, CD4-mimetic compounds and some antibodies (23). The HIV-1_AD8_ Env is efficiently expressed and processed, is well characterized with respect to antibody binding and neutralization sensitivity (Tier 2) and, among primary HIV-1 Envs, exhibits an intermediate level of triggerability by CD4 (7, 23, 73). Single, double and triple sets of lysine substitutions were introduced into the wild-type HIV-1_AD8_ Env. For example, double sets included Sets 1 + 2, 1 + 3, 2 + 3, 2 + 4, 3 + 5, etc.; triple sets included Sets 1 + 2 + 3, 2 + 3 + 4, 2 + 3 + 6, 3 + 6 + 7, etc. (Fig. 1A). In a preliminary study, a total of 24 Env variants were analyzed for protein expression and processing, ability to support entry of a pseudotyped virus, and the sensitivity of the viral pseudotype to neutralization by the 19b antibody. The 19b antibody is a poorly neutralizing antibody that recognizes the gp120 V3 loop and serves as a sensitive indicator of HIV-1 Env transitions to State-2/3 conformations (45, 71–74, 137). With a few exceptions, most of the lysine substitutions were well tolerated with respect to HIV-1_AD8_ Env processing, virus infectivity and sensitivity to 19b neutralization (data not shown). However, Envs with Set 3 + 7 and Set 3 + 6 +7 changes were poorly processed and inefficiently supported pseudovirus infection. Viruses with Set 3 + 5 changes were more sensitive than the wild-type HIV-1_AD8_ to neutralization by the 19b antibody (data not shown). Thus, while most of the introduced lysine substitutions were well tolerated, some specific combinations apparently exert undesirable effects on HIV-1_AD8_ Env conformation and function.

### Lysine-rich 2-4 R and 2-4 RED2 Envs

Based on the results of our preliminary analysis, we selected the 2-4 R Env, which contains Set 2 + 4 and R315K changes, for more detailed characterization. The ED2 set of acidic substitutions was also added to the 2-4 R Env to create the 2-4 RED2 Env (Fig. 1B). Both 2-4 R and 2-4 RED2 Envs mediated pseudovirus infection as efficiently as the wild-type HIV-1_AD8_ Env (data not shown). To evaluate Env expression, proteolytic processing and gp120-trimer association, HOS cells were transfected with plasmids expressing the wild-type HIV-1_AD8_ Env and the 2-4 R and 2-4 RED2 Envs tagged at the C-terminus with His_6_. Cell lysates were Western blotted directly (Input) or were precipitated with nickel-nitrilotriacetic (Ni-NTA) beads in the presence of BMS-806, sCD4 or the DMSO control. BMS-806 is a small-molecule HIV-1 entry inhibitor that binds gp120 and stabilizes a State-1-like Env conformation (12, 78, 138–140). The uncleaved gp160 Env precursor and mature gp120 and gp41 glycoproteins were detected in lysates of cells expressing the wild-type HIV-1_AD8_, 2-4 R and 2-4 RED2 Envs (Fig. 2A). Comparison of the gp120:gp160 ratio in the cell lysates indicates that the 2-4 R and 2-4 RED2 Envs are processed more efficiently than the wild-type HIV-1_AD8_ Env (Fig. 2A, Input lanes). In the DMSO control sample, although wild-type HIV-1_AD8_ gp41 and gp160 were precipitated by the Ni-NTA beads, little gp120 was coprecipitated (Fig. 2A, Ni-NTA lanes). Apparently, under these conditions, gp120 dissociates from the wild-type HIV-1_AD8_ Env complex. BMS-806 increased the association of the wild-type HIV-1_AD8_ gp120 with the precipitated Env complex, as previously seen (138). In the presence of sCD4, no coprecipitated gp120 was detected, presumably as a result of CD4-induced gp120 shedding (141, 142). Compared with the wild-type HIV-1_AD8_ Env, the 2-4 R gp120 was precipitated more efficiently by the Ni-NTA beads in the DMSO control lysates. The coprecipitation of the 2-4 RED2 gp120 from the DMSO-treated cell lysates by the Ni-NTA beads was even more efficient. For both 2-4 R and 2-4 RED2 Envs, the association of gp120 with the Env complex was enhanced by BMS-806 and decreased by sCD4. Thus, the Env changes in 2-4 R and 2-4 RED2 can enhance Env processing and, in detergent lysates, strengthen the association of gp120 with solubilized Env trimers. Both phenotypes were more pronounced for the 2-4 RED2 Env than for the 2-4 R Env.

**Figure 2.**
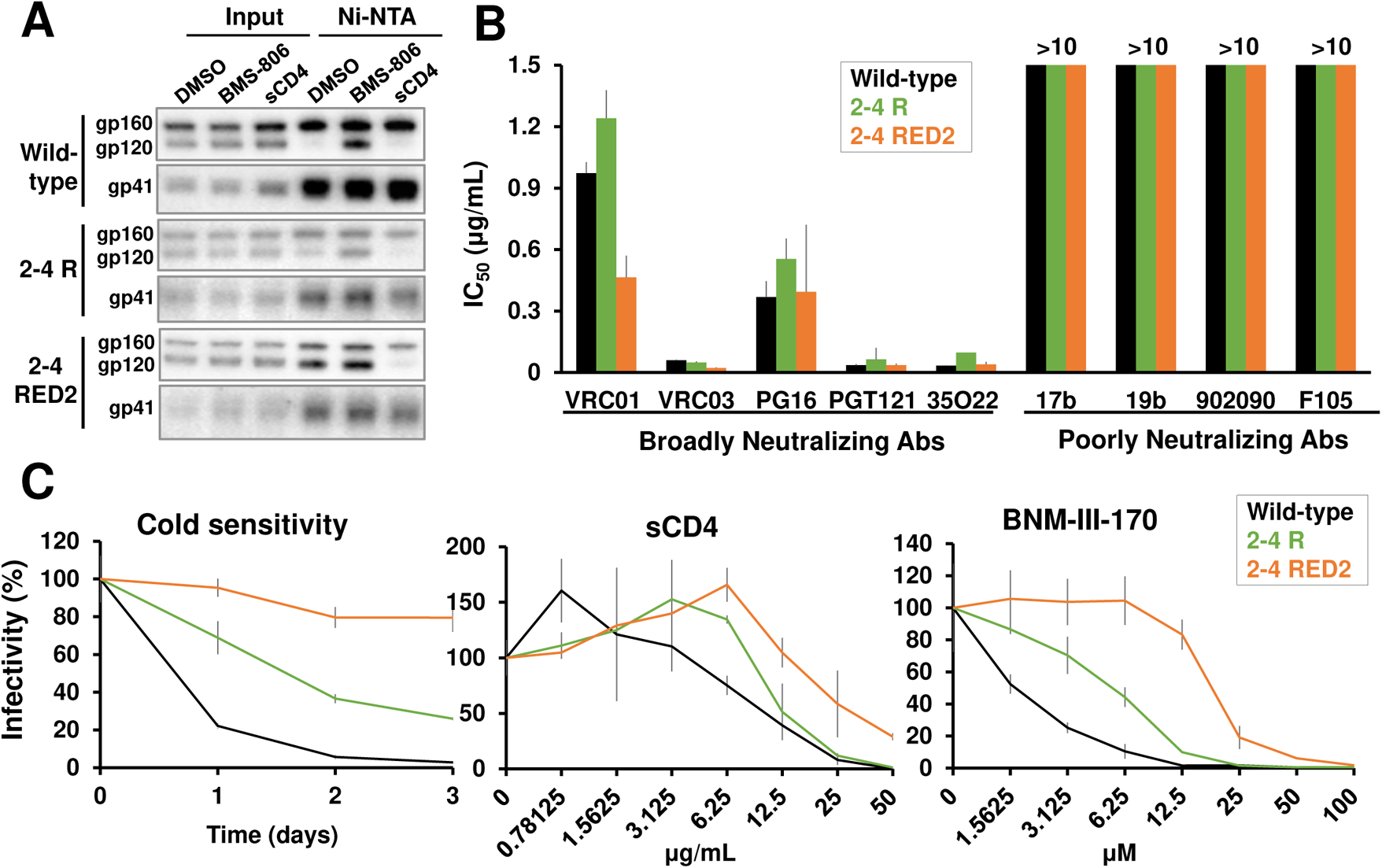
Phenotypes of the 2-4 R and 2-4 RED2 Envs. (A) HOS cells were transfected transiently with plasmids expressing His_6_-tagged wild-type HIV-1_AD8_ Env or the 2-4 R or 2-4 RED2 Env variants. Forty-eight hours later, cells were lysed; the cell lysates were incubated with Ni-NTA beads for 1.5 hr at 4°C in the presence of the DMSO control, 10 μM BMS-806 or 10 μg/mL sCD4. Total cell lysates (Input) and proteins bound to the Ni-NTA beads were Western blotted with a goat anti-gp120 antibody (upper panels) or the 4E10 anti-gp41 antibody (lower panels). (B) 293T cells were transfected with plasmids encoding the indicated Envs, HIV-1 packaging proteins and Tat, and a luciferase-expressing HIV-1 vector. Forty-eight hours later, cell supernatants were filtered (0.45 μm) and incubated with different antibodies for 1 hr at 37°C before the mixture was added to Cf2Th target cells expressing CD4 and CCR5. Forty-eight hours after infection, the target cells were lysed, and the luciferase activity was measured. The concentration of antibody required to inhibit 50% of virus infection (IC_50_) was calculated using the GraphPad Prism program. (C) Filtered cell supernatants containing recombinant viruses were incubated with sCD4 or the CD4-mimetic compound BNM-III-170 for 1 hr at 37°C. Then the mixture was added to target cells as described above. In the cold sensitivity assay, viruses were incubated on ice for the indicated times, after which the virus infectivity was measured. The results shown in A and C are representative of those obtained in at least two independent experiments. The means and standard deviations derived from two independent experiments or triplicate measurements are shown in B and C, respectively.

The sensitivity of viruses with the wild-type HIV-1_AD8_, 2-4 R and 2-4 RED2 Envs to neutralization by broadly and poorly neutralizing antibodies was examined. The broadly neutralizing antibodies (bNAbs) in our panel included VRC01 and VRC03 against the CD4-binding site of gp120 (143, 144), PG16 against a quaternary V2 epitope (145), PGT121 against a V3-glycan epitope on gp120 (146), and 35O22 against the gp120-gp41 interface (147). The poorly neutralizing antibodies included 17b against a CD4-induced epitope (148), 19b against the gp120 V3 loop (137), 902090 against a V2 gp120 epitope (149) and F105 against the CD4-binding site of gp120 (150). The 2-4 R and 2-4 RED2 viruses were neutralized by bNAbs comparably to the wild-type HIV-1_AD8_; like the wild-type HIV-1_AD8_, the 2-4 R and 2-4 RED2 viruses were resistant to poorly neutralizing antibodies (Fig. 2B).

The sensitivity of HIV-1 to inactivation by exposure to cold, sCD4 or CD4-mimetic compounds can provide an indication of Env “triggerability,” the tendency to make transitions from State 1 (23, 37, 45, 118, 120–125). Compared with the wild-type HIV-1_AD8_, the 2-4 R virus displayed slight but reproducible resistance to cold, sCD4 and BNM-III-170, a CD4-mimetic compound (151) (Fig. 2C). The 2-4 RED2 virus exhibited an even higher level of resistance to cold, sCD4 and BNM-III-170 than either the wild-type or the 2-4 R virus. These phenotypes are consistent with the stability of the State-1 Env conformation exhibiting the following rank order in these variants: 2-4 RED2 > 2-4 R > wild-type HIV-1_AD8_.

### Q114E and Q576K changes determine State 1-stabilizing phenotypes

We wished to identify the changes in 2-4 RED2 and 2-4 R responsible for the above phenotypes. Because differences among the wild-type HIV-1_AD8_, 2-4 R and 2-4 RED2 Envs were most apparent in the Ni-NTA coprecipitation and virus sensitivity experiments, we used these assays to characterize HIV-1_AD8_ Env mutants with single-residue changes corresponding to those in the 2-4 R and 2-4 RED2 Envs. Among the acidic residue substitutions found in the ED2 set, a single change, Q114E, was sufficient to recapitulate the 2-4 RED2 Env phenotypes (Fig. 3A). Similarly, a single lysine substitution originally found in Set 4, Q576K, was responsible for most of the 2-4 R Env phenotypes (Fig. 3B). Thus, Q114E or Q567K alone can enhance HIV-1_AD8_ Env processing, gp120-trimer association and virus resistance to cold, sCD4 and a CD4-mimetic compound.

**Figure 3.**
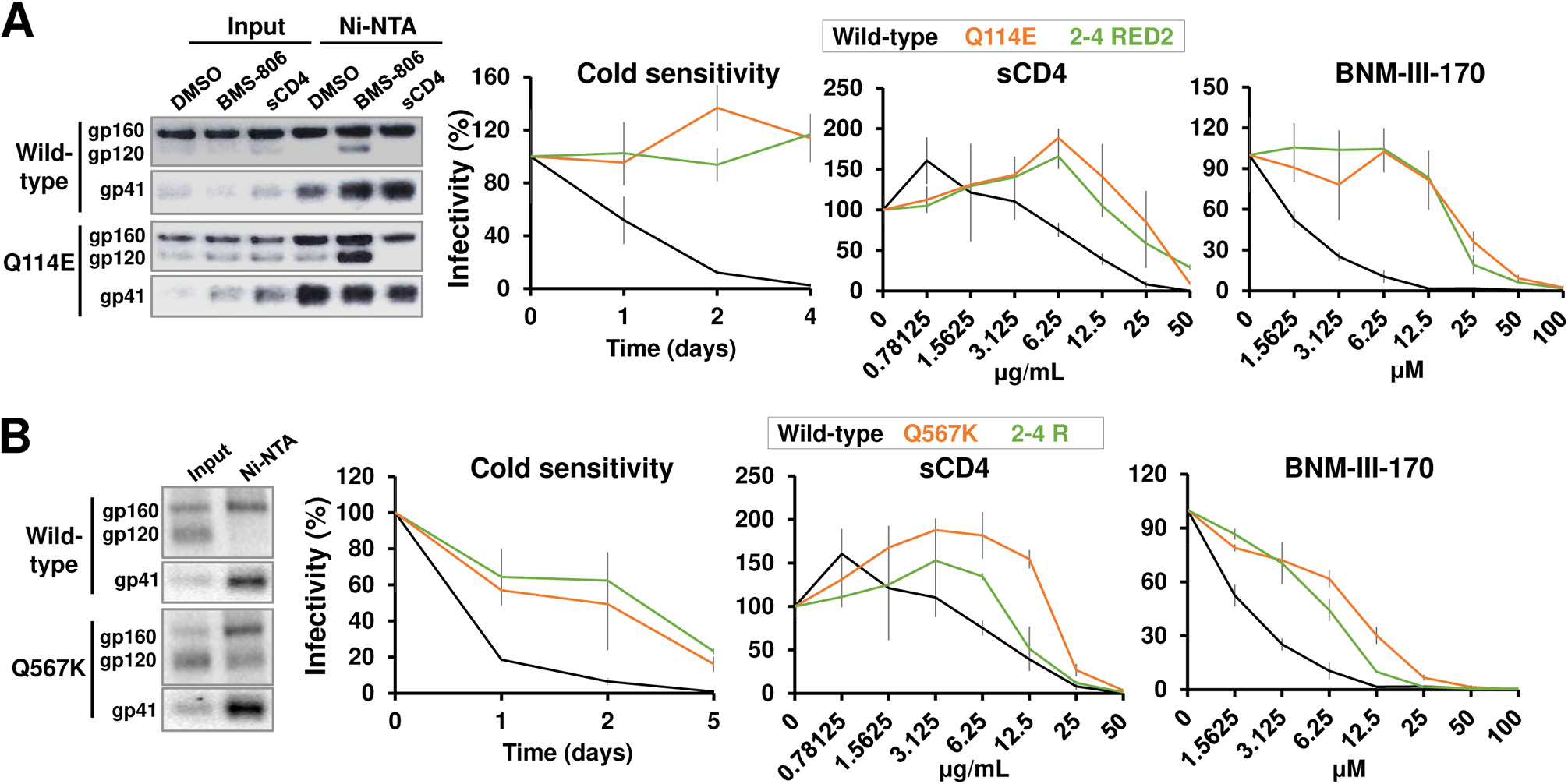
Major contributions of the Q114E and Q567K changes to the respective 2-4 RED2 and 2-4 R phenotypes. (A) The effects of the Q114E change on gp120-trimer association (left panel) and virus sensitivity to cold, sCD4 and BNM-III-170 (right panels) were measured as described in the legend to Figure 2. The sensitivities of viruses with the wild-type HIV-1_AD8_ Env and the 2-4 RED2 Env are shown for comparison. (B) The effects of the Q567K change on gp120-trimer association (left panel) and virus sensitivity to cold, sCD4 and BNM-III-170 (right panels) were measured. The sensitivities of viruses with the wild-type HIV-1_AD8_ Env and 2-4 R Env are shown for comparison. The results shown are typical of those obtained in at least two independent experiments. The right panels report the means and standard deviations derived from triplicate measurements.

Gln 114 is located in the gp120 α1 helix, part of the gp120 inner domain that faces the trimer axis and interacts with gp41 (79–82, 152–155). Gln 567 resides in the N-terminal segment of the gp41 heptad repeat 1 (HR1_N_) region, which participates in the formation of the gp41 coiled coil after CD4 binding (32–34). In the available Env trimer structures, which have been suggested to represent a State-2-like conformation (78), the HR1_N_ region is disordered or structurally heterogeneous (79–89). Although structural information on Gln 114 and Gln 567 in the context of a State-1 Env is currently unavailable, based on their approximate location near the trimer axis and the charge complementarity of the substitutions yielding similar phenotypes, we tested their functional dependence. The phenotypes of a panel of 18 single and double Q114/Q567 Env variants were characterized (Table 1). Only acidic residue substitutions at position 114 resulted in an improvement of the constellation of State-1-associated phenotypes. At position 567, lysine substitution yielded the strongest State-1-associated phenotypes, while arginine substitution exerted a more modest effect. Analysis of the double mutants yielded two insights. First, the phenotypes of the Q114E mutant were not significantly affected by changing Gln 567 to an alanine residue. Likewise, the phenotypes of the Q567K mutant were similar to those of the Q567K/Q114A double mutant. Therefore, the State-1-associated phenotypes of the Q114E and Q567K mutants are not dependent on the formation of hydrogen bonds between the side chains of residues 114 and 567. Second, the phenotypic effects of the changes in residues 114 and 567 were additive. Combination of the strongest individual changes yielded the variant, Q114E/Q567K, with the most pronounced phenotype. Both changes are found in the 2-4 RED2 Env. In summary, the Q114E and Q567K changes independently impart their individual effects on Env function and these effects are additive.

**Table 1.** Phenotypes of HIV-1_AD8_ Env variants with changes in Gln 114 and Gln 567. The phenotypes of the wild-type HIV-1_AD8_ Env and the indicated Gln 114 and Gln 567 variants were determined as in Figure 2. The values for Env processing efficiency, virus infectivity and gp120-trimer association, relative to those observed for the wild-type HIV-1_AD8_ Env, are shown. The sensitivity or resistance of viruses with the Env variants to cold, sCD4 and BNM-III-170 is reported relative to that of the wild-type HIV-1_AD8_ virus. To ensure accurate comparison of the Env variant phenotypes across multiple assays, the wild-type HIV-1_AD8_ and key Env mutants (e.g., Q114E or Q567K) were included in all assays. Phenotypes are labelled as follows: •, wild-type level; +, increase; -, decrease; R, resistant; S, sensitive; ND, not determined; NA, not applicable. For virus infectivity: 0-25 % of wild-type, - - -; 25-50 %, - -; 50-75 %, -; 75-125 %, •; >125 %, +. The data shown are representative of results obtained in at least two independent experiments.

| Env | Env Processing | Infectivity (%) | Resistance / Sensitivity compared to wild-type |  |  | gp120-trimer association |
| --- | --- | --- | --- | --- | --- | --- |
|  |  |  | Cold | sCD4 | BNM-III-170 |  |
| Wild-type | ● | ● | ● | ● | ● | ● |
| Q114A | ● | ● | ● | ● | ● | + |
| Q114D | + | ● | RR | RR | RR | ++ |
| Q114E | + | + | RR | RR | RR | ++ |
| Q114K | No | ND | ND | ND | ND | NA |
| Q114N | ● | ND | ND | ND | ND | ● |
| Q114S | ● | ND | ND | ND | ND | ● |
| Q567A | ● | + | ● | ● | ● | ● |
| Q567E | ● | - - - | ● | ● | ● | ● |
| Q567K | + | ● | R | R | R | ++ |
| Q567R | + | - | Slight R | ● | ● | + |
| Q114A Q567K | ● | ● | Slight R | R | R | ++ |
| Q114A Q567R | ● | ● | ● | ● | ● | + |
| Q114D Q567R | + | ● | R | RR | RR | ++ |
| Q114E Q567A | + | + | RR | RR | RR | ++ |
| Q114E Q567K | + | ● | RRR | RRR | RRR | ++ |
| Q114E Q567E | + | - | R | ● | R | ++ |
| Q114E Q567R | + | ● | RR | RR | RR | ++ |
| Q114K Q567E | No | ND | ND | ND | ND | NA |

We extended our mutagenesis approach to evaluate the potential of other Env residues to influence the Q114E and Q567K phenotypes. A State-1 Env structure would be most relevant to the search for interacting partners, but is currently not available. Therefore, we used the available structural models, many of which represent State-2-like Env conformations (78), to suggest candidate amino acid residues. In sgp140 SOSIP.664 trimers, the highly conserved His 72 is located ~8 Å from Gln 114 (79–81). Replacing His 72 with lysine or glutamine residues resulted in increased sensitivity to sCD4 and BNM-III-170; these phenotypes were partially relieved when these His 72 changes were combined with Q114E (Table 2). Replacing His 72 with an alanine residue resulted in a virus with neutralization sensitivity similar to that of the wild-type virus. Compared with the Q114E virus, the H72A/Q114E virus was less resistant to cold and sCD4. Thus, some changes in His 72 result in an apparent increase in Env triggerability and can influence the Q114E phenotypes.

**Table 2.**
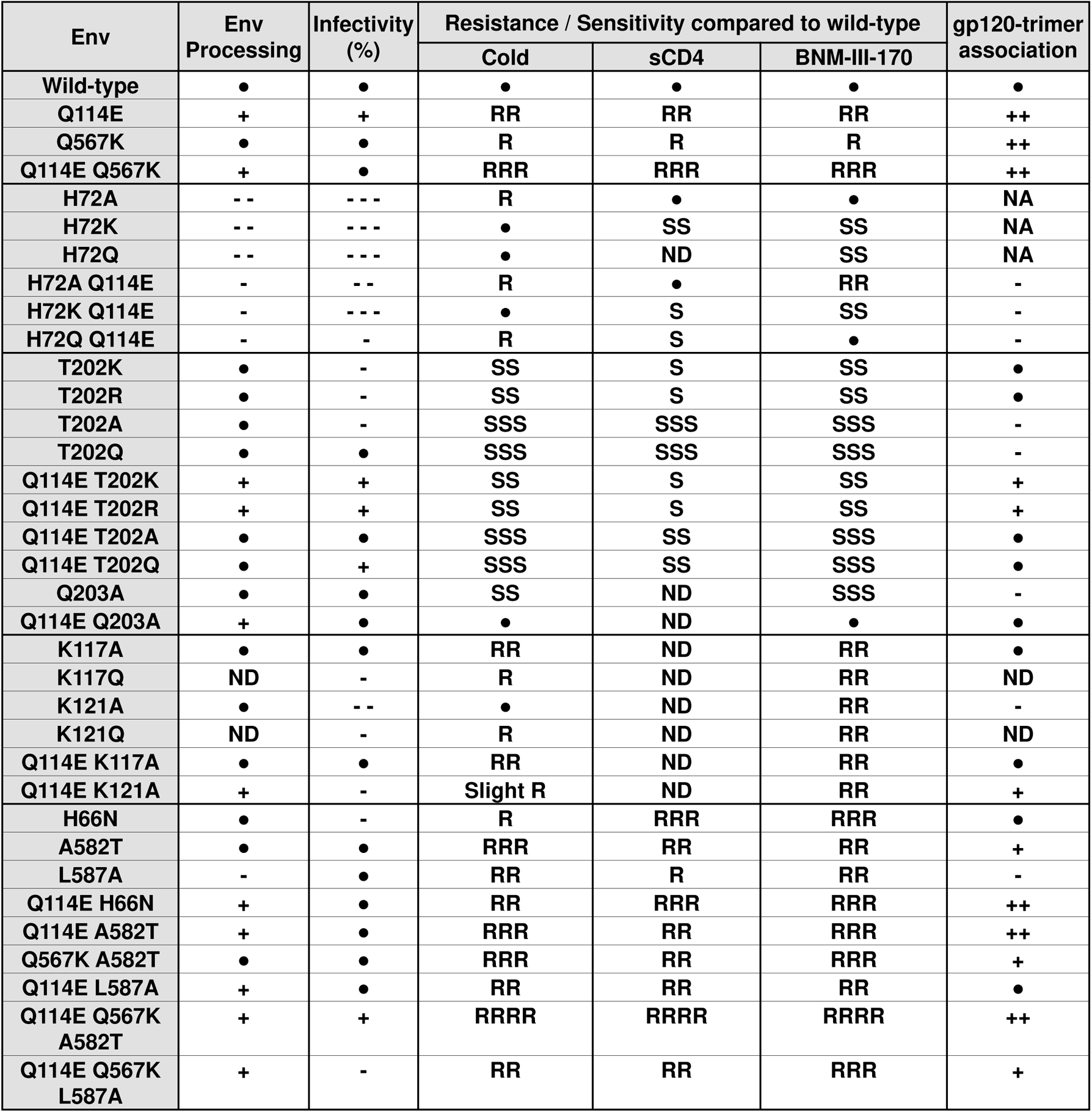
Effects of Env amino acid changes on the phenotypes of the Q114E and Q567K Env variants. The phenotypes of the wild-type and mutant HIV-1_AD8_ Envs were determined as in Figure 2. The values, relative to those of the wild-type HIV-1_AD8_ Env, are reported as described in the legend to Table 1. The data shown are representative of results obtained in at least two independent experiments.

In HIV-1/SIV_cpz_ Envs, Thr/Lys polymorphism in residue 202 often exhibits covariance with Gln/Glu polymorphism in residue 114 (136). Compared with the wild-type HIV-1_AD8_, viruses with Thr 202 replaced by alanine, lysine, arginine or glutamine residues were more sensitive to cold, BNM-III-170 and the 19b anti-V3 antibody (Table 2 and data not shown). These phenotypes, which are indicative of increased Env triggerability and State 1 destabilization, were minimally compensated by the addition of the Q114E change. Replacing the conserved Gln 203 residue with an alanine residue (Q203A in Table 2) also resulted in a State-1-destabilized phenotype, but in this case, the Q114E/Q203A mutant exhibited phenotypes close to that of the wild-type HIV-1_AD8_. Thus, the Q114E change can compensate for some but not all State 1-destabilizing changes.

In the unliganded sgp140 SOSIP.664 and PGT151-bound EnvΔCT structures (PDB: 4ZMJ and 5 FUU, respectively) (82, 86), the side chains of Gln 114, Lys 117 and Lys 121 from each Env protomer point towards the trimer axis, stacking in three layers. Interprotomer Lys 117-Lys 117 and Lys 121-Lys 121 crosslinks were formed in a crosslinking/mass spectrometry study of the sgp140 SOSIP.664 trimer, confirming the location of these residues in the trimer core in these Env structures (73). Substitution of Lys 117 or Lys 121 with an alanine or glutamine residue resulted in viruses that were more resistant to cold and BNM-III-170 than the wild-type virus (Table 2). No additive or synergistic effect was observed when the Q114E change was combined with the K117A or K121A changes. In fact, the double mutants exhibited less stable association of gp120 with solubilized Env trimers (Table 2). Thus, the effects of the Gln 114, Lys 117 and Lys 121 changes on the viral phenotypes are redundant, whereas in the detergent-solubilized Envs, the K117A and K121A changes nullify the trimer-stabilizing effects of the Q114E change. Similar phenotypic effects of the K117A and K121A changes were observed in the context of the E.2 and AE.2 HIV-1_AD8_ constructs discussed below (Table 3).

**Table 3.**
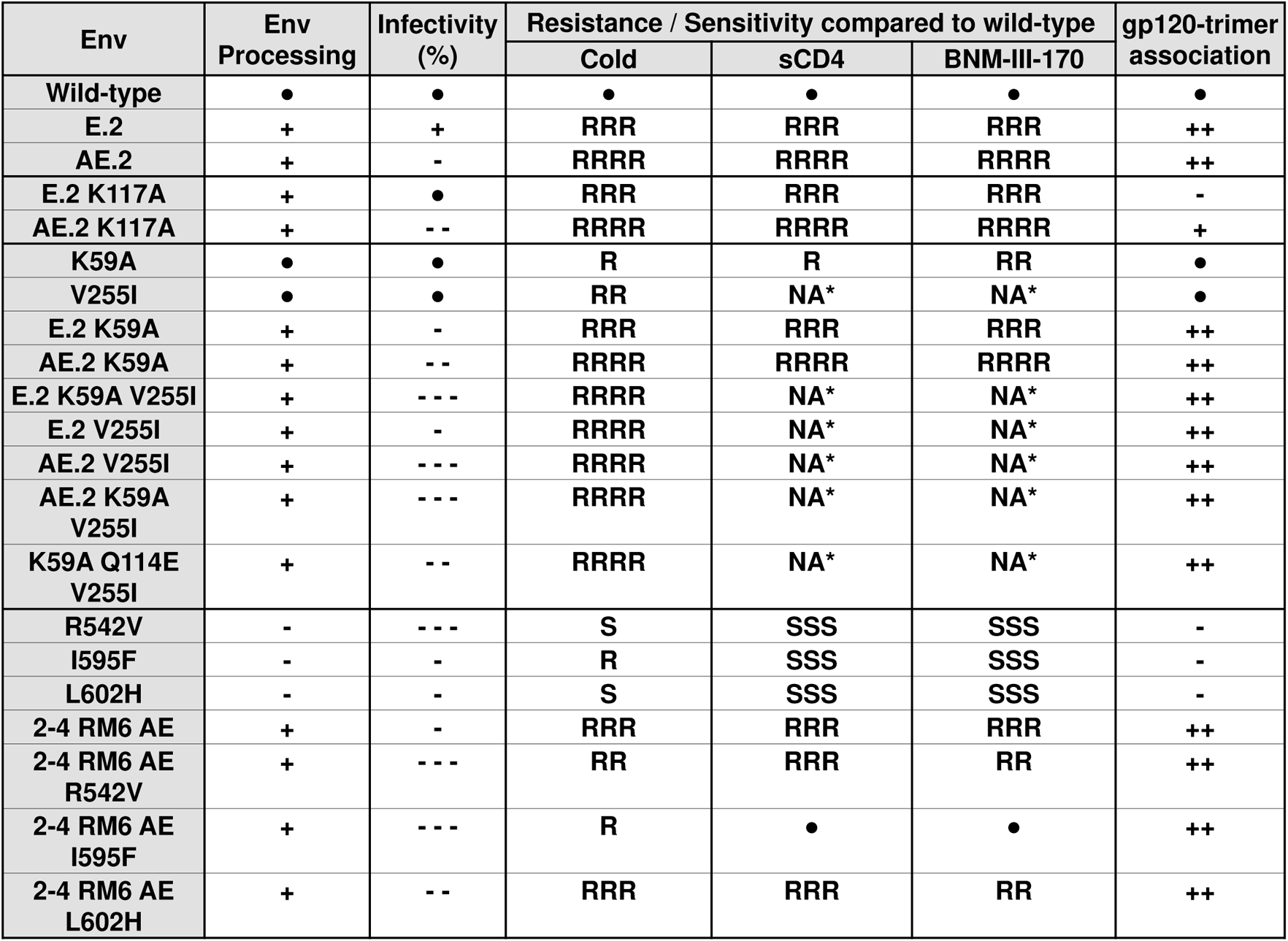
Effects of Env amino acid changes on the phenotypes of the E.2, AE.2 and 2-4 RM6 AE Env variants. The indicated amino acid changes were introduced into the HIV-1_AD8_ Env or into the E.2, AE.2 or 2-4 RM6 AE Envs. The phenotypes of these Env variants were determined as in Figure 2. The values, relative to those of the wild-type HIV-1_AD8_ Env, are reported as described in the Table 1 legend. The data shown are representative of results obtained in at least two independent experiments. *Val 255 is near the binding site for sCD4 and the CD4-mimetic compounds; therefore, the V255I change may directly decrease the binding of these Env ligands.

| Env | Env Processing | Infectivity (%) | Resistance / Sensitivity compared to wild-type |  |  | gp120-trimer association |
| --- | --- | --- | --- | --- | --- | --- |
|  |  |  | Cold | sCD4 | BNM-III-170 |  |
| Wild-type | ● | ● | ● | ● | ● | ● |
| E.2 | + | + | RRR | RRR | RRR | ++ |
| AE.2 | + | - | RRRR | RRRR | RRRR | ++ |
| E.2 K117A | + | ● | RRR | RRR | RRR | - |
| AE.2 K117A | + | -- | RRRR | RRRR | RRRR | + |
| K59A | ● | ● | R | R | RR | ● |
| V255I | ● | ● | RR | NA* | NA* | ● |
| E.2 K59A | + | - | RRR | RRR | RRR | ++ |
| AE.2 K59A | + | -- | RRRR | RRRR | RRRR | ++ |
| E.2 K59A V255I | + | --- | RRRR | NA* | NA* | ++ |
| E.2 V255I | + | - | RRRR | NA* | NA* | ++ |
| AE.2 V255I | + | --- | RRRR | NA* | NA* | ++ |
| AE.2 K59A V255I | + | --- | RRRR | NA* | NA* | ++ |
| K59A Q114E V255I | + | -- | RRRR | NA* | NA* | ++ |
| R542V | - | --- | S | SSS | SSS | - |
| I595F | - | - | R | SSS | SSS | - |
| L602H | - | - | S | SSS | SSS | - |
| 2-4 RM6 AE | + | - | RRR | RRR | RRR | ++ |
| 2-4 RM6 AE R542V | + | --- | RR | RRR | RR | ++ |
| 2-4 RM6 AE I595F | + | --- | R | ● | ● | ++ |
| 2-4 RM6 AE L602H | + | -- | RRR | RRR | RR | ++ |

As Gln 567 is disordered in most Env trimer structures, we used a low-resolution model of the uncleaved HIV-1_JR-FL_ Env (156) to suggest potential interaction partners. However, alanine substitutions in these potentially interacting HIV-1_AD8_ residues (Glu 47, Glu 83, Glu 87, Glu 91, Asp 230, Glu 492 and Glu 560) did not affect the phenotypes of the Q567K mutant virus (data not shown).

### Q114E and Q567K synergize with other State 1-stabilizing Env changes

Previous studies suggested that changes in His 66, Ala 582 and Leu 587 could enrich the State-1 HIV-1_YU2_ Env conformation through different proposed mechanisms: H66N destabilizes the CD4-bound conformation, A582T directly stabilizes the pretriggered conformation and L587A destabilizes the gp41 3-helix bundle (121, 122, 125). We confirmed that individually these changes increased HIV-1_AD8_ resistance to cold, sCD4 and BNM-III-170 (Table 2). Of the three changes, only A582T enhanced gp120-trimer association in cell lysates. Both the H66N and A582T changes synergized with the Q114E and Q567K changes in producing viral phenotypes associated with State-1 stabilization (Table 2). A combination of three changes in the Q114E/Q567K/A582T Env resulted in the most robust phenotypes.

### Crosslinkable E.2 and AE.2 Envs with enhanced State-1 stability

To generate HIV-1 Envs enriched in a pretriggered conformation and containing multiple lysine residues for crosslinking, we added two benign changes (R252K, A667K) and Q114E to the lysine-rich 2-4 R Env to create the E.2 Env construct (Fig. 1). The AE.2 Env contains, in addition, the A582T change. The A582T change was chosen because it not only resulted in viral phenotypes additive with those of Q114E and Q567K, but also increased gp120 association with the detergent-solubilized Env, a property that K117A, K121A, H66N and L587A lacked (Table 2). Both E.2 and AE.2 Env were cleaved more efficiently than the wild-type HIV-1_AD8_ Env and resisted gp120 dissociation from the solubilized Env trimer (Fig. 4A). By comparison, the wild-type Env from another primary strain, HIV-1_JR-FL_, was poorly processed and highly unstable in detergent.

**Figure 4.**
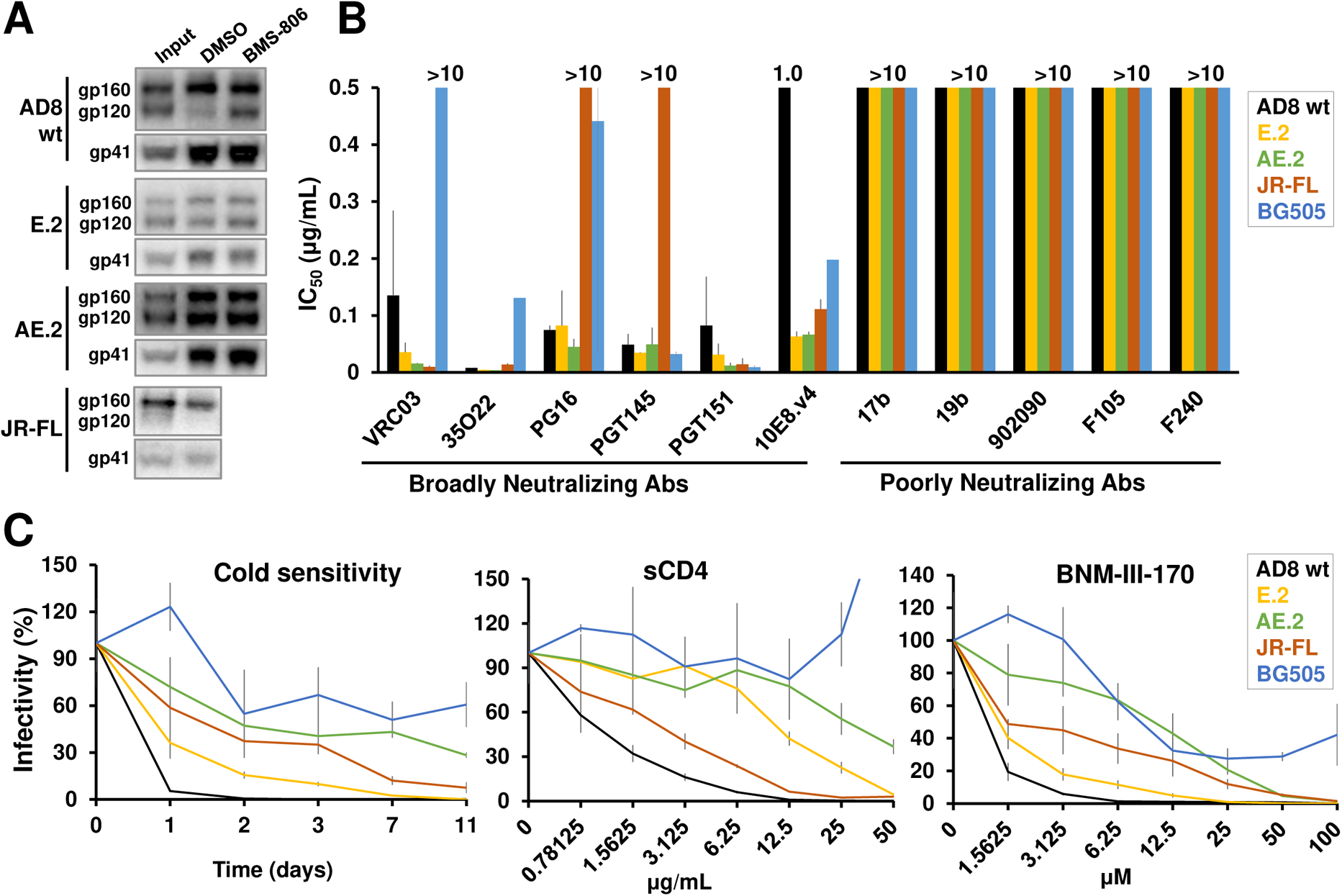
Phenotypes of the E.2 and AE.2 Envs. (A) HOS cells transiently expressing His_6_-tagged Envs (wild-type HIV-1_AD8_ Env, the E.2 Env, the AE.2 Env or the HIV-1_JR-FL_ Env) were lysed. Cell lysates were incubated with Ni-NTA beads for 1.5 hr at 4°C in the presence of the DMSO control or 10 μM BMS-806. Total cell lysates (Input) and Ni-NTA-bound proteins were Western blotted with a goat anti-gp120 antibody (upper panels) and the 4E10 anti-gp41 antibody (lower panels). (B) Recombinant luciferase-expressing viruses with the indicated Envs were incubated with antibodies for 1 hr at 37°C, after which the mixture was added to Cf2Th-CD4/CCR5 target cells. Forty-eight hours later, the target cells were lysed and the luciferase activity was measured. The IC_50_ values were calculated using the GraphPad Prism program. (C) Recombinant luciferase-expressing viruses with the indicated Envs were incubated with sCD4 or BNM-III-170 for 1 hr at 37°C before the mixture was added to Cf2Th-CD4/CCR5 target cells. Cold sensitivity was assessed by incubation of the viruses on ice for the indicated times, after which virus infectivity was measured on Cf2Th-CD4/CCR5 cells as described above. The results are representative of those obtained in at least two independent experiments. The values reported in B and C represent the means and standard deviations from at least two independent experiments or triplicate measurements, respectively.

To evaluate the functional E.2 and AE.2 Envs in more detail, we tested virus sensitivity to a panel of broadly and poorly neutralizing antibodies. In addition to the antibodies used in Figure 2, we included two bNAbs, PGT151 against the gp120-gp41 interface (157) and PGT145 against a quaternary V2 epitope (158), and the poorly neutralizing F240 antibody against gp41 (159). Envs from the Clade B HIV-1_JR-FL_ and Clade A HIV-1_BG505_ Tier 2/3 strains were included for comparison. All Env variants resisted neutralization by poorly neutralizing antibodies, as expected (Fig. 4B). Compared to the wild-type HIV-1_AD8_, HIV-1_JR-FL_ and HIV-1_BG505_, the E.2 and AE.2 viruses were just as sensitive, and even more sensitive in some cases, to neutralization by broadly neutralizing antibodies.

The sensitivity of the viruses to cold inactivation, sCD4 and the CD4-mimetic compound, BNM-III-170, is shown in Fig. 4C. Compared with the wild-type HIV-1_AD8_, the E.2 virus exhibited increased resistance to cold, sCD4 and BNM-III-170. Alteration of Glu 114 in the E.2 Env to glutamine largely reverted these phenotypes, suggesting that the Q114E change is a critical determinant of the stabilized pretriggered conformation in the E.2 Env (data not shown). The inclusion of the A582T change in the AE.2 Env further increased cold, sCD4 and BNM-III-170 resistance to the levels of the Tier 2/3 HIV-1_JR-FL_ and HIV-1_BG505_ strains. In addition, the E.2 and AE.2 viruses were more sensitive than the wild-type HIV-1_AD8_ to the State 1-preferring entry inhibitors, BMS-806 and 484 (45, 118); the AE.2 virus was more sensitive to these small-molecule inhibitors than the E.2 virus (data not shown).

In an attempt to improve the E.2 and AE.2 Envs further, we added the K59A and/or V255I changes. Lysine 59 is a highly conserved residue in the gp120 inner domain, within the disulfide loop (Layer 1) that includes His 66, discussed above. Valine 255 packs against the critical Trp 112 and Trp 427 residues in the CD4-binding Phe 43 cavity of gp120 (152); the V255I change was associated with resistance to AAR 029b, a cyclic peptide triazole inhibitor of CD4 binding (160). The K59A and V255I changes alone rendered HIV-1_AD8_ more cold-resistant, and the K59A virus was also relatively resistant to sCD4 and BNM-III-170 (Table 3). However, the K59A and V255I changes had only modest effects in the E.2 and AE.2 background on State 1-associated phenotypes, but led to significant reductions in infectivity (Table 3). These observations hint that further stabilization of State 1-associated phenotypes in the AE.2 context may be accompanied by decreases in Env function.

### Effects of State 1-destabilizing changes in different Env contexts

In the above studies, the Q114E change could revert the viral phenotypes associated with State 1 destabilization by the Q203A change but not by changes in the adjacent Thr 202 residue (Table 2). We evaluated whether an Env with multiple State 1-stabilizing changes, 2-4 RM6 AE, would better tolerate State 1 destabilization. The 2-4 RM6 AE and AE.2 Envs are identical except for the benign R252K change in the latter (Fig. 1). The 2-4 RM6 AE virus is resistant to cold, sCD4 and BNM-III-170 and exhibits a strong gp120-trimer association in detergent (Table 3). We individually introduced the R542V, I595F and L602H changes into the wild-type HIV-1_AD8_ Env or the 2-4 RM6 AE Env. These gp41 changes rendered HIV-1 more sensitive to the nonpeptidic inhibitory compound RPR103611, which suggested that they might destabilize the pretriggered (State-1) Env conformation (161). In agreement with this hypothesis, the R542V and L602H viruses exhibited increased sensitivity to cold, sCD4 and BNM-III-170 relative to HIV-1_AD8_ (Table 3). The I595F virus was sensitive to sCD4 and BNM-III-170 as well as to the 19b anti-V3 antibody, but was slightly more resistant to cold inactivation than HIV-1_AD8_. Interestingly, the increased sensitivity to cold, sCD4, BNM-III-170 and 19b associated with these gp41 changes was not evident in the 2-4 RM6 AE background. Thus, the State 1-stabilizing changes in 2-4 RM6 AE apparently resist the State 1-destabilizing effects of the R542V, I595F and L602H changes in the gp41 ectodomain.

### Correlations among key Env phenotypes

To understand the relationships among key Env phenotypes and to visualize the effects of specific amino acid changes on the progression of successive generations of Env mutants, we plotted the relative levels of resistance to cold, BNM-III-170 and gp120-trimer dissociation for all characterized Env variants (Fig. 5). Virus resistance to cold inactivation reflects the stability of the functional Env trimer on virions and is independent of the binding of an Env ligand. Virus resistance to the CD4-mimetic compound generally correlates with resistance to sCD4 (122,154, 162). Of interest, there exists a strong correlation between virus resistance to the CD4-mimetic compound and to cold (Fig. 5). Beginning with the wild-type HIV-1_AD8_ Env, Envs incorporating additive State 1-stabilizing changes displayed upward shifts towards highly resistant phenotypes, comparable to those of the HIV-1_JR-FL_ and HIV-1_BG505_ Envs. Envs with State 1-destabilizing changes grouped together in the lower left quadrant.

**Figure 5.**
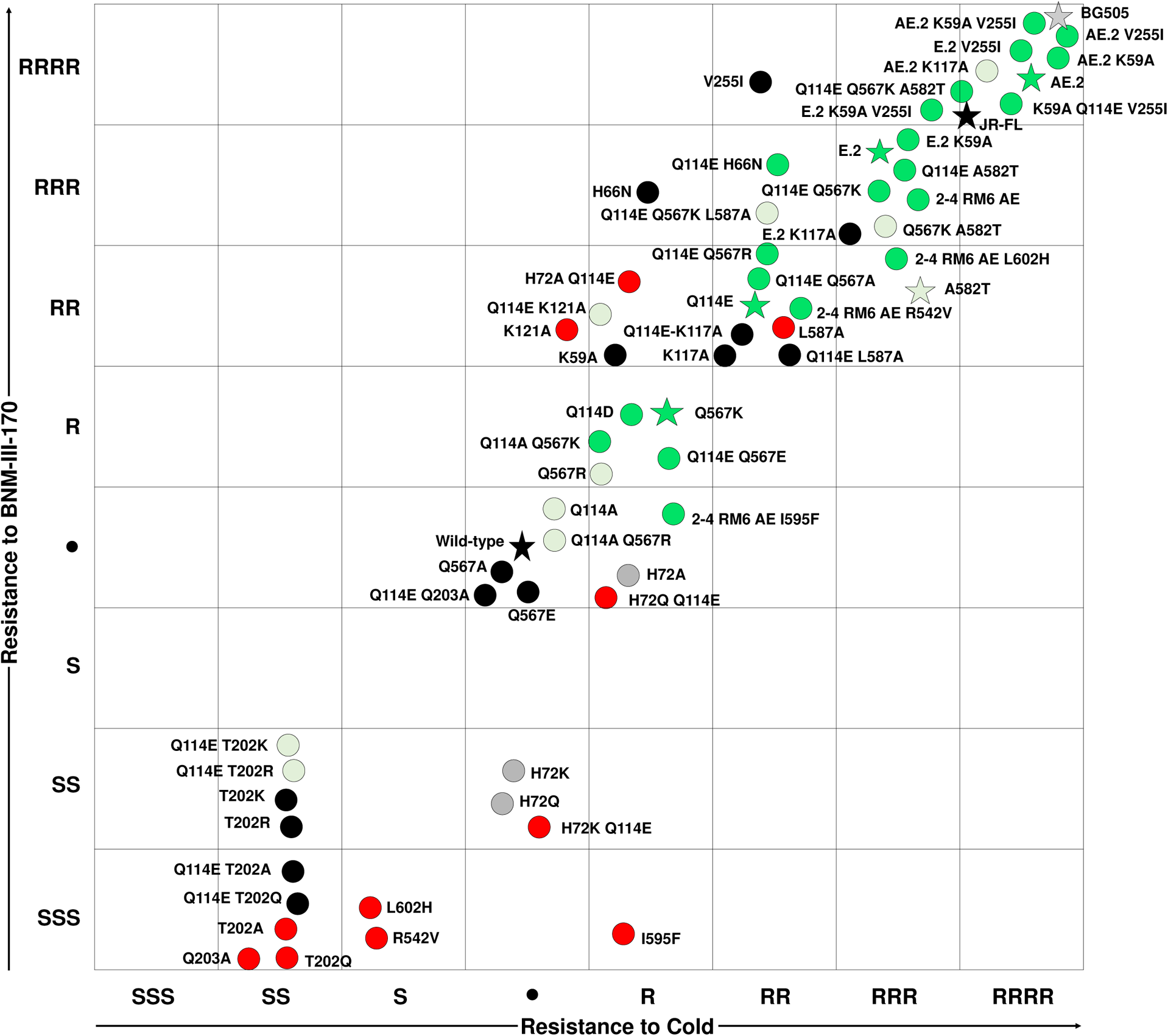
Correlations among key Env phenotypes. The plot shows the relative level of resistance to the CD4-mimetic compound BNM-III-170 versus the relative level of cold resistance for the HIV-1 Env variants tested in this study. The levels of resistance are scored as described in the legends to Tables 1–3: •, wild-type level; R, resistant; S, sensitive. Key Env variants are designated with stars. Envs are colored according to their relative gp120-trimer association level, as measured by Ni-NTA coprecipitation of gp120 with the His_6_-tagged gp41 glycoprotein: black, wild-type level; gray, not determined or not applicable; light green, +; green, ++; red, -. The V255I change is located near the binding site for CD4-mimetic compounds and may directly affect Env interaction with BNM-III-170. Note that the E.2 and AE.2 Envs exhibit resistance to cold and BNM-III-170 comparable to those of the HIV-1_JR-FL_ and HIV-1_BG505_ Envs, but also display better gp160 processing and a tighter association of gp120 with the Env trimer solubilized in detergent.

Env variants that exhibited a higher level of gp120-trimer association in detergent, relative to that of the wild-type HIV-1_AD8_ Env, are colored green in Figure 5. The skewed distribution of these Env variants in the upper right quadrant indicates that a tighter association of gp120 with the solubilized Env trimer is related to virus resistance to cold and BNM-III-170, phenotypes associated with State 1 stabilization.

Note that several Env variants, including the natural HIV-1_JR-FL_ and HIV-1_BG505_ Envs, achieve virus resistance to cold and BNM-III-170 without increasing gp120-trimer association in detergent-solubilized Envs. Therefore, increasing gp120-trimer association is not the only means of achieving a more stable pretriggered (State-1) Env conformation.

### Crosslinking efficiency of the wild-type AD8, E.2 and AE.2 Envs

The lysine-rich E.2 and AE.2 Envs are expected to crosslink more efficiently than the wild-type HIV-1_AD8_ Env with lysine-reactive crosslinkers like DTSSP and glutaraldehyde. DTSSP has a spacer arm of 12 Å, whereas, because of its tendency to polymerize, glutaraldehyde forms crosslinks of more variable lengths (163). Both DTSSP and glutaraldehyde crosslinked the E.2 and AE.2 Envs more efficiently than the wild-type AD8 Env (Fig. 6A). For example, after treatment with 5 mM glutaraldehyde, the E.2 and AE.2 Envs crosslinked into gel-stable trimers, whereas the wild-type HIV-1_AD8_ Env mostly formed monomers and dimers. Apparently, a greater number of lysine residues accessible to the crosslinkers exist on the surface of the E.2 and AE.2 Env trimers compared with the wild-type HIV-1_AD8_ Env.

**Figure 6.**
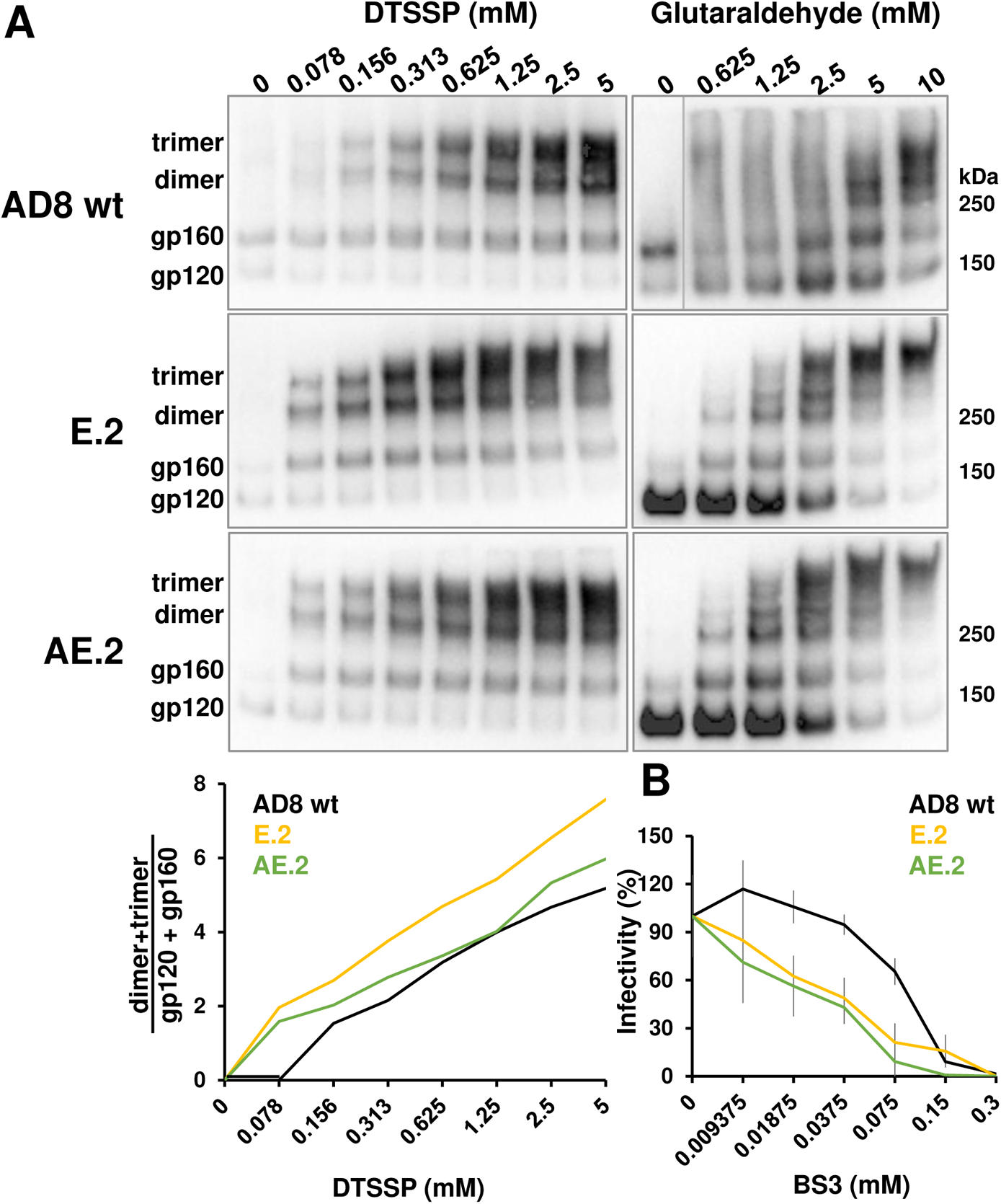
Crosslinking of the wild-type HIV-1_AD8_, E.2 and AE.2 Envs. (A) VLPs consisting of the HIV-1 Gag-mCherry fusion protein and the wild-type HIV-1_AD8_ Env, the E.2 Env or the AE.2 Env were incubated with different concentrations of the DTSSP or glutaraldehyde crosslinkers. After quenching the reactions, VLPs were pelleted and lysed. The VLP proteins were analyzed by reducing or non-reducing PAGE, respectively, followed by Western blotting. The ratio of gel-stable (dimers + trimers):(gp120 + gp160) provides an indication of interprotomer crosslinking by DTSSP. (B) Luciferase-expressing viruses pseudotyped with the wild-type HIV-1_AD8_, E.2 or AE.2 Envs were incubated with the BS3 crosslinker. After quenching the reaction, the viruses were added to Cf2Th-CD4/CCR5 cells. Luciferase activity in the target cells was measured 48 hours later. The results shown in A and B are representative of those obtained in at least two independent experiments. The values reported in B represent the means and standard deviations derived from triplicate measurements.

We also examined the relative sensitivity of the functional wild-type HIV-1_AD8_, E.2 and AE.2 Envs to BS3, another lysine-specific crosslinker with spacer arms of 12 Å. The infectivity of viruses pseudotyped with the E.2 and AE.2 Envs was inhibited by BS3 at three- to four-fold lower concentrations than those required for inhibition of viruses with the wild-type HIV-1_AD8_ Env (Fig. 6B). These results suggest that BS3 crosslinks occur more efficiently on the E.2 and AE.2 Envs than on the wild-type HIV-1_AD8_ Env, leading to a loss of infectivity at lower BS3 concentrations.

## DISCUSSION

Despite more than three decades of intense research, an effective HIV-1 vaccine remains elusive. The metastability and multiple conformational states of the HIV-1 Env create challenges for the generation of broadly neutralizing antibodies, either following vaccination or during natural HIV-1 infection. In Env-expressing cells, both uncleaved and cleaved (mature) Envs are present on the cell surface. A significant fraction of the uncleaved Env bypasses the conventional Golgi secretory pathway to traffic to the cell surface; these Envs differ from mature Envs in glycan processing, conformation and recognition by antibodies (7). Uncleaved Envs may function as a decoy to the host immune system and divert antibody responses away from the mature Envs. The pretriggered (State-1) conformation of the mature virion Env of primary HIV-1 strains is the target for most broadly neutralizing antibodies (12, 37, 38, 45). This native conformation, however, is unstable and can transition into more open State 2/3 conformations that are able to be recognized by poorly neutralizing antibodies. Therefore, it is of significant interest to devise methods to lock Env in its native State-1 conformation by means that resist perturbation during Env purification, characterization and immunization.

Here, we tackled the challenges posed by HIV-1 Env conformational flexibility in two ways. First, we used polymorphisms in naturally occurring HIV-1 strains to guide the introduction of extra lysine and acidic amino acid residues in the HIV-1_AD8_ Env. Chemical crosslinkers that couple lysine or acidic residues on proteins under physiological conditions are available (132–135). During the iterative process employed to identify HIV-1_AD8_ Envs that are potentially more susceptible to crosslinking, we required that the Env variants exhibit efficient processing, subunit association, and the ability to support virus entry. Some of the functional HIV-1_AD8_ Env variants developed by this approach contain up to 11 extra lysine residues (33 per Env trimer) and up to 7 extra acidic residues (21 per Env trimer). Using DTSSP or glutaraldehyde as crosslinking agents, two Env variants, E.2 and AE.2, were shown to form interprotomer crosslinks more efficiently than the wild-type HIV-1_AD8_ Env. The infectivity of viruses with these Envs was inactivated more efficiently than that of viruses with the wild-type HIV-1_AD8_ Env by another lysine-specific crosslinker, BS3. These assays document the accessibility of some of the additional lysine residues introduced into the E.2 and AE.2 Envs. Chemical crosslinking can enrich the representation of labile native conformations in Env preparations for structural analysis or immunogenicity studies. Crosslinking/mass spectrometry can provide distance constraints between Env residues that can be used to validate available structural models or to derive new models (132–135). A previous study utilized crosslinking/mass spectrometry to detect differences between soluble and membrane-bound Envs (73). The inclusion of the 2-4 RED2, E.2 and AE.2 Envs in future crosslinking/mass spectrometry studies should increase the number of distance constraints and thereby improve our ability to discriminate among alternative structural models.

The second strategy employed in our approach was to screen the Env variants for function and viral phenotypes associated with stabilization of a State-1 Env conformation. For this purpose, we evaluated viral resistance to cold, sCD4 and the CD4-mimetic compound BNM-III-170. Cold inactivation reflects the resistance of the functional HIV-1 Env trimer to the detrimental effects of ice formation at near-freezing temperatures (164–166). The sensitivity of HIV-1 variants to cold inactivation is related to the intrinsic reactivity or triggerability of Env; Envs that more readily make the transition from State 1 to downstream conformations are invariably cold-sensitive (23, 121, 122). HIV-1 sensitivity to sCD4 and BNM-III-170 inhibition is a function of Env triggerability (23, 45, 118, 119, 125); because we generally avoided changes to the highly conserved and well-defined BNM-III-170 binding site on gp120 (151, 162), most of the observed differences in virus sensitivity to this CD4-mimetic compound reflect changes in the ability of Env to negotiate transitions from a State-1 conformation. Our study documents the strong correlation between HIV-1 resistance to cold and resistance to BNM-III-170. This screening strategy identified two changes, Q114E in gp120 and Q567K in gp41, that individually increased the resistance of the HIV-1_AD8_ Env to inactivation by cold, sCD4 and BNM-III-170. These viral phenotypes were additively enhanced by combining the Q114E and Q567K changes in Env variants, such as the lysine-rich E.2 and AE.2 Envs. Cold, sCD4 and BNM-III-170 resistance were further increased by the inclusion in the AE.2 Env of the A582T gp41 change, which previously was shown to stabilize a pretriggered Env conformation (123, 125). The functional E.2 and AE.2 Envs exhibit an antigenic profile consistent with a State-1 conformation, conferring virus sensitivity to broadly neutralizing antibodies that target quaternary epitopes (PG16, PGT145, PGT151, 35O22) and resistance to poorly neutralizing antibodies (17b, 19b, 902090, F105, F240). Viruses with the E.2 and AE.2 Envs were inhibited efficiently by BMS-806, a small molecule that exhibits some preference for a State-1 Env conformation (12, 78, 138–140). Two unanticipated beneficial phenotypes associated with the E.2 and AE.2 Envs are more efficient Env processing and greater stability of solubilized Env trimers. HIV-1 Env cleavage has been suggested to contribute to the stability of the State-1 conformation (138, 139, 167–173). As uncleaved HIV-1 Envs sample multiple conformations, including those reactive with poorly neutralizing antibodies, achieving a high level of gp120-gp41 processing may be important for an effective vaccine immunogen. The E.2 and AE.2 Envs achieve levels of State 1-associated phenotypes comparable to those of the Tier 2/3 HIV-1_JR-FL_ and HIV-1_BG505_, but notably, are processed much more efficiently. In addition, relative to these Envs and the wild-type HIV-1_AD8_ Env, the E.2 and AE.2 Envs solubilized in detergent exhibit much greater gp120 association with the Env trimer. The Q114E, Q567K and A582T changes individually strengthen the non-covalent association of gp120 with the solubilized Env trimers, a property that will assist purification and characterization. Of interest, the Q567K change was included in a combination of Env changes that were reported to stabilize HIV-1 Env trimers in different contexts (174–176). In our panel of HIV-1 Env variants, enhancement of Env trimer stability in detergent was strongly correlated with virus resistance to cold and BNM-III-170, State 1-associated phenotypes. We note that the binding of the State 1-preferring compound, BMS-806, also stabilizes gp120-trimer association (138). In future studies, the ability of Q114E, Q567K and A582T changes to enhance Env cleavage efficiency, gp120-trimer association and State-1 stabilization in other HIV-1 strains will be explored.

We identified other changes (K59A, K117A, K121A) that individually yielded Env phenotypes consistent with State-1 stabilization. These and previously identified State 1-stabilizing changes (H66N, L587A) (121, 122, 125) were tested in combination with the Q114E and/or Q567K changes in various Env backgrounds. In no case did we observe an additive improvement in viral phenotypes associated with State 1 stabilization, and several of these combinations resulted in attenuated virus replication or gp120-trimer dissociation in detergent. It is not surprising that, as State 1 stability is increased, virus replication diminishes as the activation barriers governing State 1-to-State 2 transitions increase. However, the validation of State 1 stabilization would be less straightforward for replication-incompetent Envs, given current uncertainties about a State-1 Env structure. Therefore, we deferred investigation of these potentially State-1-stabilizing changes until better assays to characterize the conformations of nonfunctional Envs are established.

Changes in gp41 (I559P, L555P) that are intended to prevent the formation of the HR1 coiled coil have been used to stabilize soluble gp140 trimers (70, 88, 177). However, introduction of these changes in combination with the major State 1-stabilizing changes (Q114E/I559P, Q114E/Q567K/I559P and Q114E/Q567K/L555P) resulted in Envs that were not processed (data not shown).

We also considered another gp41 change, Q658E, that has been shown to stabilize sgp140 SOSIP.664 trimers (178). Introduction of the Q658E change into the wild-type HIV-1_AD8_ Env resulted in increased virus sensitivity to cold, sCD4, BNM-III-170 and the 19b antibody (data not shown). These phenotypes are consistent with those reported in other HIV-1 strains (178) and, as they suggest a lower occupancy of State 1, we did not evaluate the Q658E change in combination with the Q114E and Q567K changes.

Although a State-1 Env structure is currently unknown, mapping the Env residues identified in this study on available Env trimer models can provide some insights. Figure 7 shows the locations of Env residues in which changes resulted in increases or decreases in State 1-associated phenotypes on a PGT151-bound HIV-1_JR-FL_ EnvΔCT trimer (PDB: 5FUU) (82). The binding of the PGT151 antibody induces a State-2 conformation that is asymmetric, with two antibody Fabs bound to the Env trimer (78, 82). We chose this structure because, unlike most HIV-1 trimer structures, the HR1_N_ region containing Gln 567 is resolved; however, in keeping with the asymmetry of the PGT151-bound Env trimer, the positions of the Gln 567 residues differ among the three Env protomers. Gln 567, Gln 114, and Ala 582 are close to the trimer axis in the EnvΔCT structure (Fig. 7A). The C_α_-C_α_ distances between Gln 114 and Gln 567 residues vary from 11.6 to 15.2 Å and the side chains of these residues do not apparently interact in this Env conformation. Gln 114 is stacked above Lys 117 and Lys 121, the side chains of which project towards the Env trimer axis (Fig. 7B, right panel). Although a precise structural explanation for the observed State 1-stabilizing phenotypes will require more data, the implicated residues are positioned near intersubunit or interprotomer junctions and therefore could potentially modulate trimer opening. For example, electrostatic repulsion among Lys 117 residues that destabilizes the Env trimer could be mitigated by their conversion to alanine residues or by replacing Gln114 with acidic residues.

**Figure 7.**
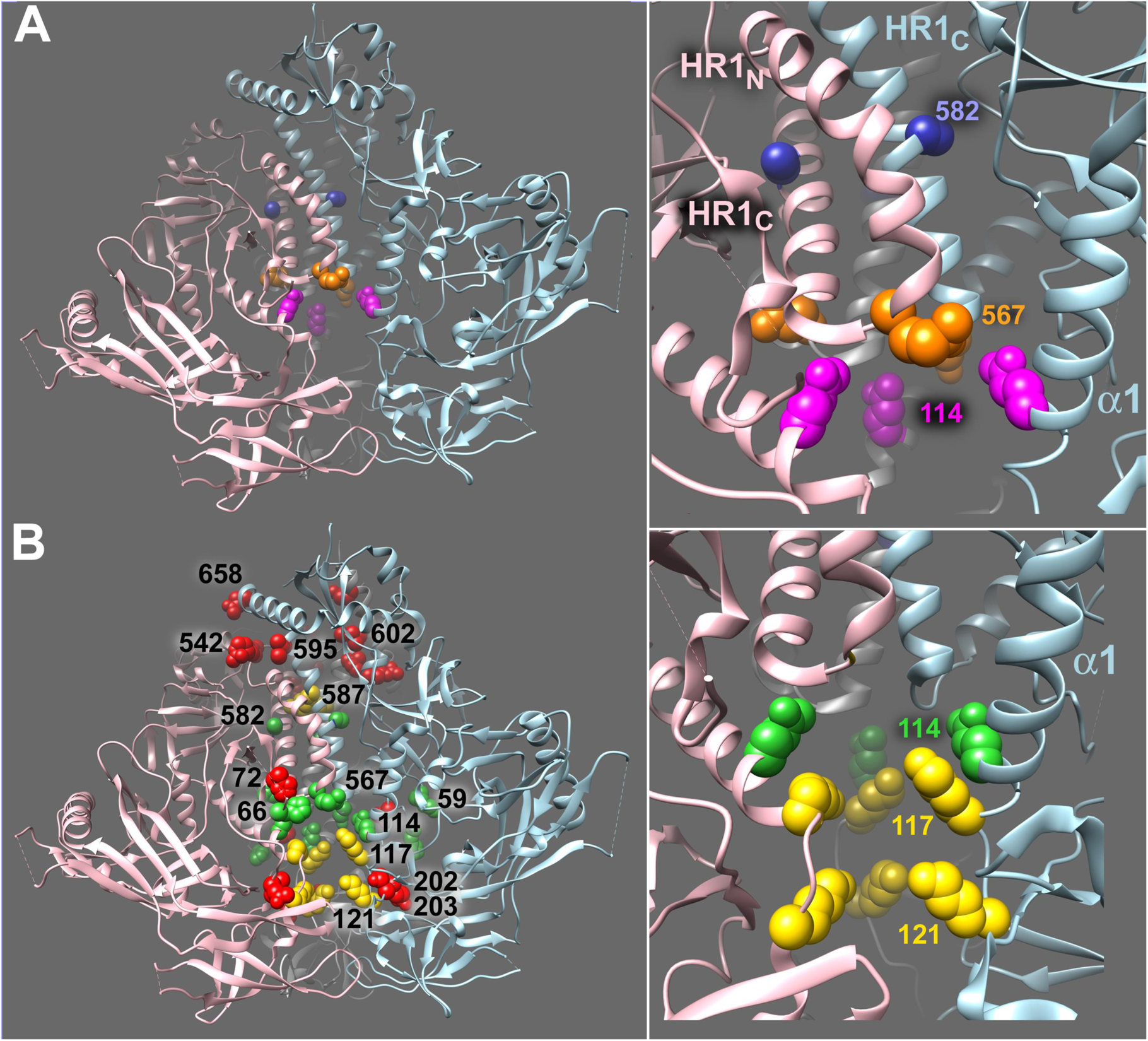
Location of Env residues in a structural model of an HIV-1 Env trimer. Env residues studied herein are depicted as CPK spheres in a PGT151-bound HIV-1_JR-FL_ Env ΔCT trimer (PDB: 5FUU) (82). The binding of two PGT151 Fabs introduces asymmetry into the Env trimer. In this depiction, the PGT151 Fabs have been removed from the structure. The individual Env protomers are colored pink, light blue and gray. In this orientation, the gp120 subunits are at the bottom and gp41 subunits at the top of the figures. (A) Env residues (Gln 114 (magenta), Gln 567 (orange) and Ala 582 (blue)) associated with State 1-stabilizing changes are shown. The distances between the C_α_ atoms of Gln 114 and Gln 567 residues in this asymmetric trimer structure are 11.6, 13.1 and 15.2 Å. The HR1_N_ regions of the three Env protomers differ in conformation. (B) Env residues (Lys 59, His 66, Gln 114, Gln 567 and Ala 582) associated with State 1-stabilizing changes are colored green. Env residues (His 72, Thr 202, Gln 203, Arg 542, Ile 595, Leu 602 and Gln 658) associated with State 1-destabilizing changes are colored red. Changes in the residues (Lys 117, Lys 121 and Leu 587) colored yellow resulted in Envs that were resistant to cold and a CD4-mimetic compound, but were subject to gp120 dissociation from the Env trimer solubilized in detergent. The right panel shows the side-chain stacking of residues Gln 114, Lys 117 and Lys 121 near the Env trimer axis.

The State 1-destabilizing changes identified in this study (red residues in Fig. 7B, left panel) are less localized than the State 1-stabilizing changes (green and yellow residues in Fig. 7B, left panel). This is consistent with the expectation that a metastable structure can be disrupted by a diverse set of changes, whereas a more limited and strategically placed set of changes is required to strengthen the structure. In this study, we provide an example of how State 1-stabilizing changes in Env can counter the phenotypic effects of State 1-destabilizing alterations, even when these changes involve amino acid residues very distant on current Env trimer structures.

In a related paper, we report the ability of the State 1-stabilizing changes identified herein to counter the phenotypic consequences of disruption of the gp41 MPER. Although further work will be required to understand fully the mechanisms underlying these observations, the ability of the Q114E, Q567K and A582T changes to counteract the disruptive effects of distant changes suggests that they may have significant utility in preserving pretriggered Env conformations in multiple circumstances.

## MATERIALS AND METHODS

### Env glycoprotein constructs

The HIV-1_AD8_ and HIV-1_JR-FL_ Envs were coexpressed with the Rev protein in the pSVIIIenv expression vector, using the natural HIV-1 *env* and *rev* sequences (23). The Asp 718 (Kpn I)-BamHI fragment of HIV-1_AD8_ *env* was cloned into the corresponding sites of the pSVIIIenv plasmid expressing the HIV-1_HxBc2_ Env and Rev. The initial single, double and triple sets of lysine substitutions shown in Fig. 1A were introduced into the HIV-1_AD8_ Env lacking an epitope tag. A carboxy-terminal GGHHHHHH (His_6_) epitope tag was added to the Env variants shown in Fig. 1B and derivatives thereof. The mutations were introduced by site-directed PCR mutagenesis using Pfu Ultra II polymerase (Agilent Technologies), according to the manufacturer’s protocol. The plasmid expressing the HIV-1_BG505_ Env (BG505.W6M.ENV.C2) was obtained through the NIH HIV Reagent Program and was contributed by Dr. Julie Overbaugh.

### Cell lines

293T cells (ATCC) and HOS cells (ATCC) were grown in Dulbecco’s Modified Eagle’s Medium/Nutrient Mixture F12 (DMEM-F12) supplemented with 10% fetal bovine serum (FBS) and 100 μg/ml of penicillin-streptomycin. A549 cells expressing HIV-1 Envs with Gag-mCherry fusion proteins were grown in DMEM-F12 medium supplemented with 10% FBS, 1X Pen-Strep, 1X L-glutamine and 0.2% Amphotericin B. Cf2Th cells stably expressing the human CD4 and CCR5 coreceptors for HIV-1 were grown in the same medium supplemented with 0.4 mg/ml of G418 and 0.2 mg/ml of hygromycin. All cell culture reagents are from Life Technologies.

### Env processing and gp120-trimer association in Ni-NTA precipitation assay

HOS cells were cotransfected with a Rev/Env-encoding pSVIIIenv plasmid and a Tat-encoding plasmid at a 1:0.125 ratio using the Effectene transfection reagent (Qiagen). At 48 h after transfection, HOS cells were washed with 1X PBS and lysed in 100 mM (NH_4_)_2_SO_4_, 20 mM Tris-HCl, pH 8, 300 mM NaCl and 1.5% Cymal-5 (Anatrace) containing DMSO, 10 μM BMS-806 or 10 μg/mL soluble CD4-Ig. Lysates were clarified and aliquots were saved as the input samples. The remaining lysates were incubated with nickel-nitriloacetic acid (Ni-NTA) beads (Qiagen) for 1.5 h at 4°C. The beads were gently pelleted and washed 3 times with room temperature washing buffer (100 mM (NH_4_)_2_SO_4_, 20 mM Tris-HCl, pH 8, 1 M NaCl and 0.5% Cymal-5). The beads were then boiled in LDS sample buffer, and the proteins analyzed by Western blotting using 1:2,000 goat anti-gp120 polyclonal antibody (Thermo Fisher Scientific) and 1:2,000 HRP-conjugated rabbit anti-goat IgG (Thermo Fisher Scientific) or 4E10 anti-gp41 antibody (Polymun) and 1:2,000 HRP-conjugated goat anti-human IgG (Santa Cruz).

### Virus infectivity, neutralization and cold sensitivity

Single-round virus infection assays were used to measure the ability of the Env variants to support virus entry, as described previously (23). Briefly, 293T cells were cotransfected with the Rev/Env-encoding pSVIIIenv plasmid; a Tat-encoding plasmid; the pCMV HIV-1 Gag-Pol packaging construct; and a plasmid containing the luciferase-expressing HIV-1 vector at a weight ratio of 1:0.125:1:3 using a standard calcium phosphate transfection protocol. At 48 h after transfection, virus-containing supernatants were collected, filtered through a 0.45-μm membrane, and incubated with soluble CD4, BNM-III-170 or antibody for 1 h at 37°C. The mixture was then added to Cf2Th-CD4/CCR5 cells, which were cultured at 37°C/5% CO_2_. To enhance infection by recombinant viruses with the HIV-1_BG505_ Env, virus-antibody mixtures were spinoculated with target cells at 1800 rpm for one hour at room temperature and then incubated for one more hour before additional medium was added. Luciferase activity in the Cf2Th-CD4/CCR5 target cells was measured 48 h later. To measure cold sensitivity, the viruses were incubated on ice for various lengths of time prior to measuring their infectivity. To measure the sensitivity of virus infectivity to crosslinking, the viruses were incubated with BS3 (Thermo Fisher Scientific) for 15 minutes at room temperature; the reaction was quenched with 15 mM Tris-HCl, pH 8.0 for 10 minutes, and the mixture was then added to the target cells.

### Crosslinking of Envs on virus-like particles (VLPs)

A549 cells inducibly expressing virus-like particles (VLPs) consisting of the HIV-1 Gag-mCherry fusion protein and the wild-type HIV-1_AD8_ Env have been previously described (7, 138). The D1253 A549-Gag/Env cell line expressing VLPs with wild-type HIV-1_AD8_ Env was selected by FACS sorting for Gag-positive and PGT145-positive cells. The D1555.042321.sort A549-Gag/E.2 Env cells and the D1553.042321.sort A549-Gag/AE.2 Env cells inducibly expressing VLPs with the E.2 and AE.2 Envs, respectively, were established similarly. After FACS sorting, these cells were >90% dual-positive for Gag expression (KC567 antibody-positive) and Env expression (PGT145 antibody-positive).

An equivalent number of cells from the three cell lines described above were seeded and the expression of Gag-mCherry/Env VLPs was induced with 2 µg/ml doxycycline. Forty-eight to seventy-two hours later, supernatants containing VLPs were centrifuged at low speed to remove cell debris and then filtered (0.45 µm). Clarified supernatants were centrifuged at 100,000 x g for one hour at 4°C. VLP pellets were resuspended in 1X PBS, aliquoted and incubated with different concentrations of either DTSSP (Thermo Fisher Scientific) or glutaraldehyde crosslinkers. The crosslinking reaction with DTSSP was carried out for 30 minutes at room temperature, after which the reaction was quenched with 100 mM Tris-HCl, pH 8.0 for 10 minutes at room temperature. Glutaraldehyde crosslinking was carried out for 5 minutes at room temperature, after which the reaction was quenched with 50 mM glycine for 10 minutes at room temperature. VLPs were then pelleted at 20,000 x g for 30 minutes at 4°C. VLP pellets were resuspended in 1X PBS/LDS, boiled and Western blotted with a goat anti-gp120 antibody, as described above. The intensity of the gp120, gp160, dimer and trimer bands was quantified using the BioRad Image Lab program.

## ACKNOWLEDGMENTS

We thank Ms. Elizabeth Carpelan for manuscript preparation. Antibodies against HIV-1 were kindly supplied by Dennis Burton (Scripps), Peter Kwong and John Mascola (Vaccine Research Center NIH), Barton Haynes (Duke University), Hermann Katinger (Polymun), James Robinson (Tulane University), and Marshall Posner (Mount Sinai Medical Center). We thank the NIH HIV Reagent Program for providing additional reagents.

This work was supported by grants from the National Institutes of Health (grants AI145547, AI124982 and AI150471, including the Basic Science Core of the University of Alabama Center for AIDS Research (AI27767)) and by a gift from the late William F. McCarty-Cooper.

